# TRAWLING: a Transcriptome Reference Aware of spLIciNG events

**DOI:** 10.1101/2021.12.03.471115

**Authors:** Noemi Di Nanni, Alejandro Reyes, Daniel Ho, Robert Ihry, Audrey Kauffmann, Eric Y. Durand, Antoine de Weck

**Author notes:** To whom correspondence should be addressed Contact: Antoine de Weck, Noemi Di Nanni. Competing Interests Statement: During their involvement related to this reported work, all authors were employees and shareholders of Novartis.

## Abstract

Alternative splicing is critical for human gene expression regulation and plays an important role in multiple human diseases. In this context, RNA sequencing has emerged as powerful approach to detect alternative splicing events.

In parallel, fast alignment-free methods have emerged as a viable alternative to quantify gene and transcript level abundance from RNAseq data. However, the ability to detect differential splicing events is dependent on the annotation of the transcript reference provided by the user.

Here, we introduce a new reference transcriptome aware of splicing events, TRAWLING, which simplifies the detection of aberrant splicing events in a fast and simple way. In addition, we evaluate the performances and the benefits of aligning transcriptome data to TRAWLING using three different RNA sequencing datasets: whole transcriptome sequencing, single cell RNA sequencing and Digital RNA with pertUrbation of Genes.

Collectively, our comprehensive evaluation underlines the value of using TRAWLING in transcriptomic data analysis.

**Availability and implementation:** Our code is available at https://github.com/Novartis/TRAWLING

## Introduction

The journey from gene to protein is complex and consists mainly of two processes: transcription and translation. During transcription, genes’ DNA is transcribed into messenger RNA. Coupled to transcription, introns are removed from RNAs by a process called RNA splicing, which also enables cells to selectively exclude alternative exons from the final matured mRNA. The resulting transcripts can generate diverse protein isoforms (Clancy, 2008; Le et al., 2015) through translation by ribosomes.

Recent studies estimated that up to 90-95% of human genes with multiple exons are alternatively spliced, highlighting the importance of pre-mRNA splicing in encoding distinct transcript isoforms and maintaining organism functionality (Le et al., 2015; Baralle et al., 2017; Wei et al. 2021). Therefore, any disruptions of normal splicing patterns can potentially affect the generation of functional proteins and cause a wide range of human diseases, such as cancer, muscular dystrophy, and other common human disorders (Scotti et al., 2016; Garcia-Blanco et al., 2004). These findings suggest that the detection of abnormal regulation of alternative splicing is of critical importance to identify diagnostic biomarkers and potential therapeutic targets (Le et al., 2015; Wu et al., 2019; Bessa et al., 2020; Zhang et al. 2020).

In the past few years, RNA sequencing (RNA-Seq) has emerged as a powerful technology to analyze transcriptomes and to study alternative splicing at both bulk and single-cell levels (Arzaluz-Luque et al., 2018; Liang et al., 2021; Liu et al., 2021). To detect isoform-switching events, it is crucial to quantifying transcriptomes at the transcript level, since the use of gene counts for statistical analysis can mask transcript-level dynamics (Zhang et al., 2017; Booeshaghi et al., 2021). In addition, the ability to detect differential isoform usage is dependent on the transcript reference annotation provided by the user (Zhang et al., 2017).

In this study, we present TRAWLING, a Transcriptome Reference AWare of spLicING events. TRAWLING simplifies the identification of splicing events from RNA-seq data in a simple and fast way, while leveraging the suite of tools developed for alignment-free methods (e.g. Kallisto (Bray et al., 2016), Salmon (Patro et al., 2017), Alevin (Srivastava et al., 2019)). In addition, it allows the aggregation of read counts based on the donor and acceptor splice motifs. As proof of concept, we evaluated TRAWLING on three different transcriptome datasets: whole transcriptome sequencing (Shuai et al. (2019)), single cell RNA sequencing (Johnston et al. (2020)) and Digital RNA with pertUrbation of Genes (Li J et al., (2021)).

## Materials and Methods

### TRAWLING

TRAWLING is built starting from gtf annotation file (Figure 1, Supplementary Figure 1) as input, and outputs a transformed gtf file. The transformation follows five main steps (Figure 2):

**Figure 1.**
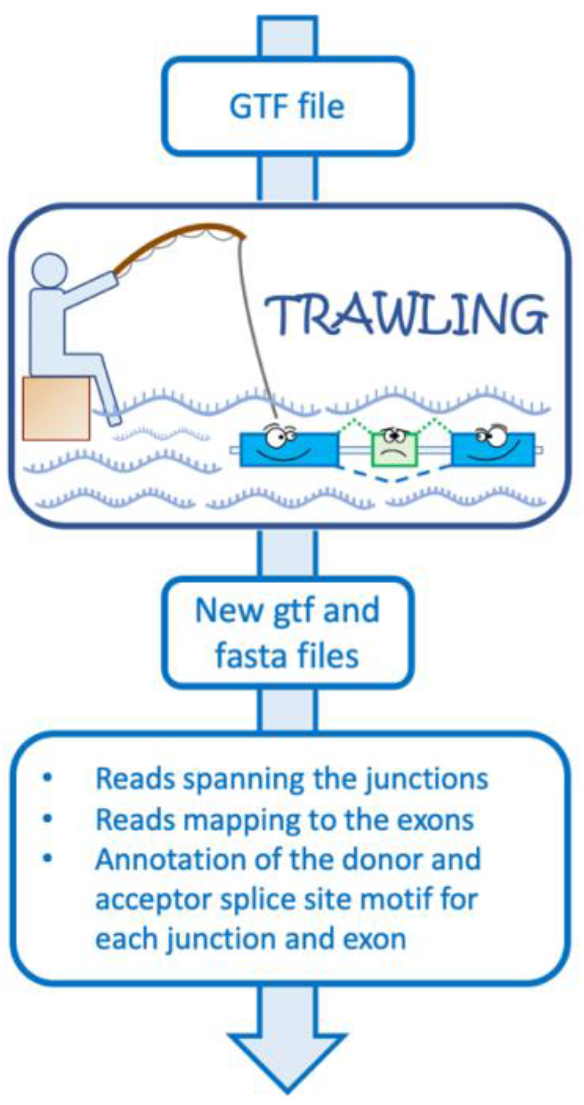
TRAWLING. TRAWLING is built from a gtf annotation file (e.g. GENCODE). It allows the easy detection of splicing events and estimation of exon and junction specific (pseudo)-mapping. In addition, it allows the aggregation of read counts based on donor and acceptor splice site motifs.

**Figure 2.**
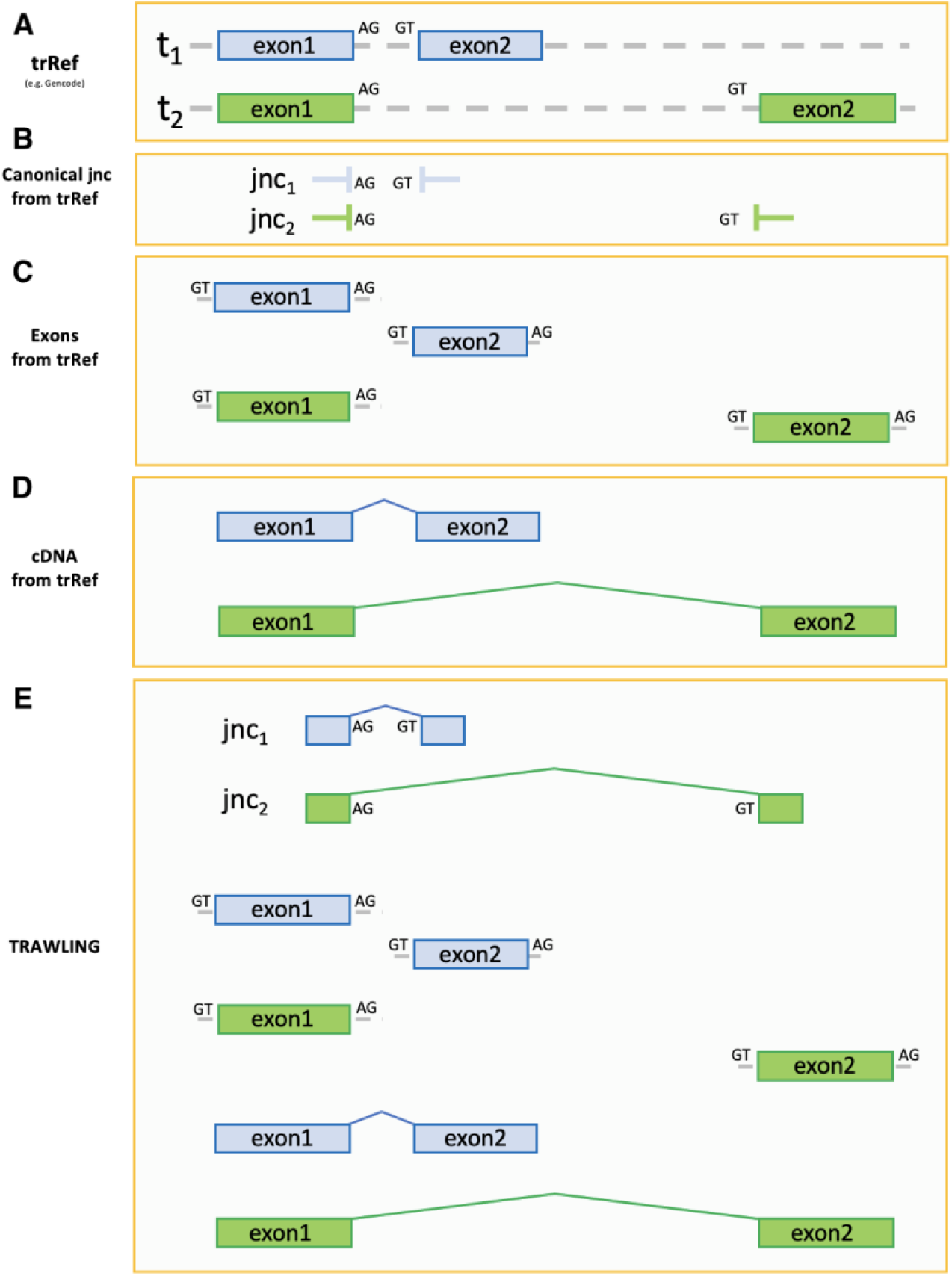
Main ideas of TRAWLING. In this conceptual example, (A) for a given transcript T two isoforms (t_1_ and t_2_) are annotated in the transcriptome reference (trRef). (B) The exonic sequences flanking the annotated introns are captured in order to record junction specific reads. (C) The exon sequences are extracted and considered as transcripts themselves. (D) The cDNA sequences of each transcript are extracted. (E) TRAWLING is built including the sequences obtained at the (B)-(C)-(D) steps. Additionally, each TRAWLING “transcript” in steps (B)-(C) is annotated with its respective donor and acceptor splice site motifs; in particular, 4 nucleotides within the exons and 10 nucleotides within the introns are annotated for both donor and acceptor splice motifs.

**Step1. Junction annotation**: annotation of the sequences at the exon-exon junctions (Figure 2B) for each transcript. In particular, *n* nucleotides within each exon are included respective to the starting and ending position of the intron. The value of *n* should be chosen based on the reads’ length, in a way that allows to capture the information about how many reads are spanning a given junction. Typically, n is set to the read length minus one read, or minus up to 10% of the read length. As a result, each junction is annotated in the output gtf file as a transcript with two exons.

**Step2. Exon annotation**: annotation of each exon’s sequence and consider it as a transcript itself (Figure 2C). The exons shorter than *n* nucleotide are discarded to avoid incorrect alignment but are included in the next step within the cDNA (Supplementary Figure 1).

**Step3. Splice motifs annotation**: the donor and acceptor splice motifs are annotated for each junction and exon. In particular, 4 nucleotides within the exons and 10 nucleotides within the introns are annotated for both splice motifs. The information is annotated in a bed file format and the sequences are extracted using bedtools (Aaron et al., 2010).

**Step4. cDNA annotation**: annotation of the cDNA for each isoform (Figure 2D).

**Step5. Reference construction and creation**: the information collected in the previous steps is used to build TRAWLING (Figure 2E) and is annotated in a gtf file format. Lastly, the transcript reference fasta file is obtained using the TRAWLING’s GTF file and the gffread tool (Pertea et al., 2020) (Supplementary Figure 2).

### Data

For the following results TRAWLING was built from GENCODE gtf file and was evaluated on whole transcript and 3’ RNA sequencing datasets.

#### Transcriptome references

The hg38 human reference genome and the gtf annotation file were downloaded from GENCODE (Ratajczak et al., 2021). They were used to get the cDNA transcript reference fasta file using gffread tool and to build TRAWLING (Supplementary Figure 1-2). Two versions of the cDNA transcript reference were created. The first is based on full transcripts and used on full transcripts RNAseq results. The second restricts the referenced transcripts to the last 400 nucleotides of each annotated transcripts only. These were then used to pseudo-align 3’ RNA sequencing datasets. Lastly, since TRAWLING depends on read length and since it was evaluated on three difference datasets, three versions of TRAWLING were built, as reported below.

#### 3’ RNA sequencing datasets

##### DRUG-seq data

We used a publicly available DRUG-seq dataset (GEO accession GSM5357052) presented in Li J et al., (2021). The dataset consists of an osteosarcoma cell line treated with 14 compounds (Table 1), each with an 8-point dose response (3.2 nM to 10 uM). In addition, 8 DMSO wells were included in the experiment.

**Table 1.** Compounds included In the DRUG-seq dataset. Table depicts compound name, identifier, target, from Li J et al. 2021.

| Compound | NSN identifier | Target |
| --- | --- | --- |
| Triptolide | ND09-QY33 | ERCC3i |
| CHIR118637 | CA-90-VK59 | GSK3Bi |
| Cmp_341 | JC-68-AL90 | JAK2i |
| Fedratinib | MB-03-IE36 | JAK2i |
| NVS-SM2 | JC-43-ZO95 | SNRPCmod |
| CPI-203 | PA-33-CK45 | BRD4i |
| Brutasol | KA-73-NB69 | NRF2i |
| Homoharringtonine | QC-05-UB63 | EIF4Ei |
| BTdCPU | SE-15-AV21 | EIF2AK1a |
| AZD8055 | NV-67-DX31 | MTORi |
| Cmp_334 | DB-85-YA47 | PI3K |
| Dilazep | NA-37-JQ34 | SLC29A2i |
| BLU9931 | VA-76-OV33 | FGFR4 |
| (S)-Crizotinib | LD-22-SA99 | ALK |

TRAWLING was built by considering the last 400 nucleotides of cDNA of each transcript and by setting the value of *n* to 45, since the reads are 51 nucleotide long. FastQ reads were pseudoaligned using Kallisto (Bray et al., 2016).

Kallisto-bustool was then used to pseudoalign the reads to TRAWLING (*y*) as well as, for comparison, to the last 400 nucleotide of cDNA of each transcript obtained from GENCODE gtf file (*x)*. To evaluate if the alignment to *x* could lead to the loss of aligned reads, the coefficient of determination (R^2^) was calculated between *x* and *y*, using the read counts value of each gene-compound pairs. Z-scores were calculated for each treatment to identify splicing events showing a statistically significant difference when compared to the control compound (DMSO).

##### scRNA-seq data

To also evaluate TRAWLING on scRNASeq data we used the data presented in Johnston et al. (2020) (GEO accession GSM4317810). The dataset consists of 6000 cells derived from the bone marrow of a patient with acute myeloid leukemia (AML) sequenced using the Illumina NovaSeq 6000 protocol.

Again, TRAWLING was built by considering the last 400 nucleotide of cDNA of each isoform and by setting the value of *n* to 135, since the reads are 150 nucleotide long. FastQ reads were aligned using Kallisto. Kallisto-bustool was used to pseudoalign the reads to TRAWLING (*y*) and to the last 400 nucleotide of cDNA of each transcript obtained from GENCODE gtf file (*x*). R^2^ was calculated between *x* and *y* using the read counts value of each gene-cell pairs. The resulting cell-by-gene matrices were pre-processed and analysed using Seurat (Stuart et al., Cell, 2019), in particular to perform data normalization, scaling, variable gene selection, PCA, clustering, UMAP calculation and to identify cell cluster maker genes. The identified marker genes were used to assign a cell type to each cluster. Based on the markers’ expression, cell clusters n.3 and n.8 displayed signatures typical of T (Dahl et al., 1992) and B-cells (Schmidt et al., 2012) respectively (Supplementary figure 3). The remaining cell clusters were labeled as a ‘malignant cells’ cluster based on the CD34’s expression. Z-scores were used to detect splicing events showing a statistically significant difference between the malignant cells cluster and the B/T cells cluster.

**Figure 3.**
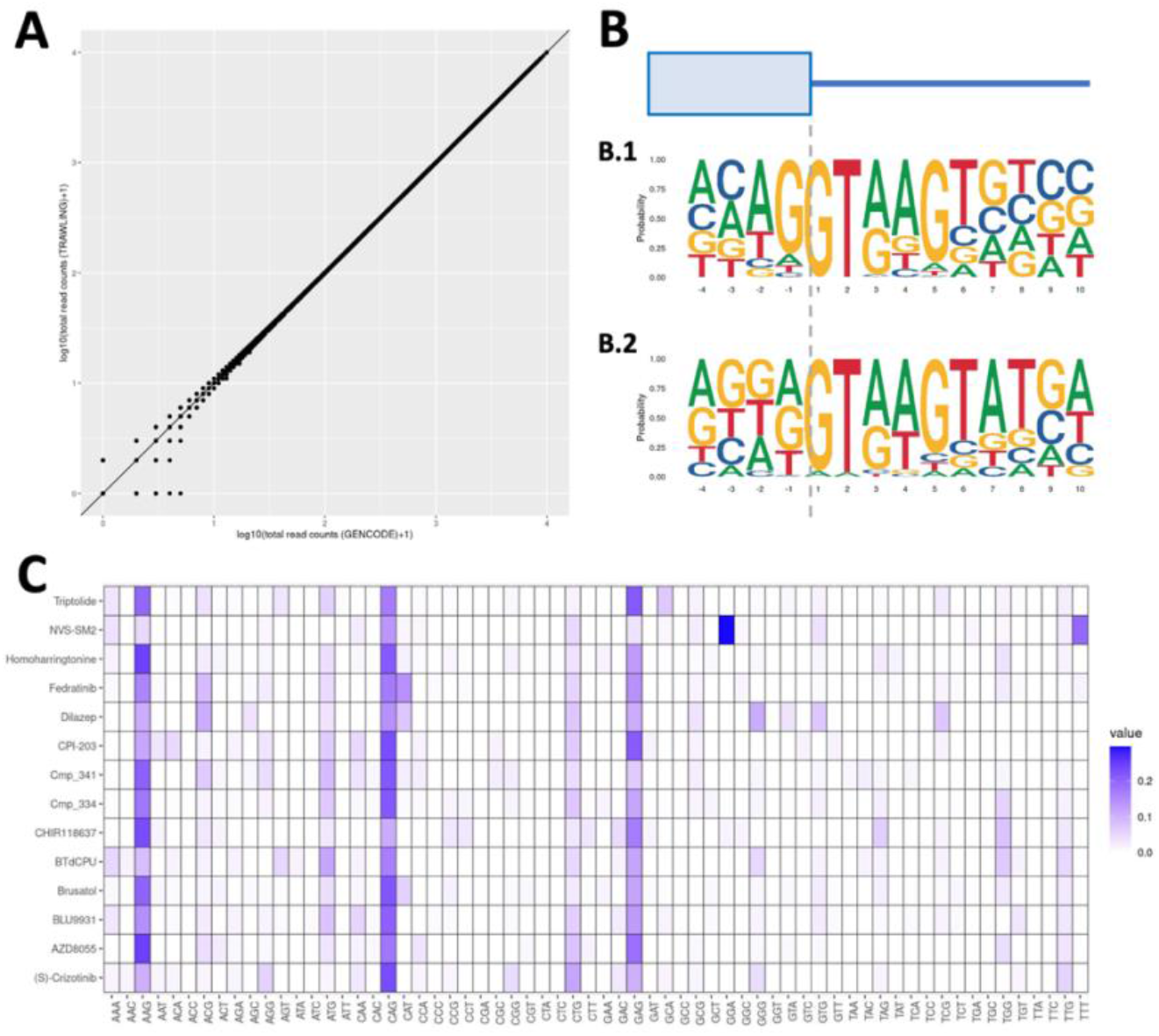
DRUG-Seq data analysis. (A) Scatter plot of gene expression levels for each compound aligned to the last 400 nucleotide of cDNA of each transcript (x-axis) and to TRAWLING (y-axis). (B) NVS-SM2 sequence selectively alters exon splicing as previously shown (*Palacino, J*., *et al. Nat Chem Biol. 2015)* (B.1) Splicing signal motif at 5’ss from (B.1) 6259 and (B.2) 38 splicing events. The former are splicing events showing a non-statistically significant difference between DMSO and NVS-SM2, contrary to the latter. (C) Heatmap showing for each compound (row) the enrichment for each motif at the 5’ss (last 3 nucleotide within the exon). The plot is generated by only including splicing events which displayed a statistically significant difference between DMSO and the considered compounds, when used at a dosage of 10uM.

#### Whole transcript RNA-seq dataset

We used two publicly available bulk RNAseq datasets (GEO accession number GSM3938668 and GSM3938669) presented in Shuai et al. (2019). The RNA-seq datasets consist of chronic lymphocytic leukaemia cell lines with (MUT, GSM3938668) or without (WT, GSM3938669) an A>C somatic mutation at the third base of the U1 snRNA.

TRAWLING was built by setting the *n* value equals to 70, since the reads are 78 nucleotide long. FastQ files were pseudoaligned using Kallisto to TRAWLING (*y*) and to the cDNA of each transcript obtained from GENCODE gtf file (*x*). R^2^ was calculated between *x* and *y*, using the read counts value of each gene-cell pairs. Z-scores was calculated to identify splicing events showing a statistically significant difference between the MUT and WT samples and vice-versa.

## Results

TRAWLING is a transcriptome reference that easily allows the detection and quantification of alternative splicing events from RNA-seq data. As a motivating factor we demonstrate its ability to aggregate read counts based on donor/acceptor splice motifs. We then present TRAWLING’s performance in the general problems of aligning RNA-seq reads to a transcriptome reference and in detecting splicing events. The performances were evaluated on three different datasets in which we expected to detect splicing events. In particular, the DRUG-seq experiment included a splicing modulator compound among 14 compounds to treat an osteosarcoma cell line. For the scRNA-seq dataset, we selected an AML dataset because the contribution of alternative splicing to gene dysregulation in AML patients has already been demonstrated. (Rivera et al. 2021, de Necochea-Campion et al. 2016, Dlamini et al. 2020). Lastly, for the bulk RNA-seq example, we used a dataset where the cell lines are known to have U1 mutations which are expected to influence splicing in a splice site motif dependent manner.

### DRUG-seq data analysis

TRAWLING successfully recapitulated gene level counts compared to what was obtained by aligning to just the last 400 nucleotides of cDNA of each transcript (Figure 3A, R^2^ equals to 0.999999).

Furthermore, we leveraged TRAWLING’s ability to aggregate read counts based on donor/acceptor splice motifs. This information could prove useful in the discovery and understanding of compounds’ mechanism of action. As proof of concept, we focused on NVS-SM2, a splicing modulator compound that selectively alter exon splicing (Palacino et al., Nat Chem Biol 2015). We detected splicing events by looking for statistically significant difference in read-counts between NVS-SM2 and DMSO. In total, 38 events were identified and by aggregating their read counts at the 5 splice site (5’ss), NVS-SM2-responsive exons showed to be enriched for a GGA motif in the last three exonic nucleotides (as previously shown in Palacino et al., Nat Chem Biol 2015) (Figure 3B).

The same analysis was applied to each compound included in the DRUG-seq dataset to check whether other compounds induced differential splicing of exons and whether responsive exons were enriched for a specific motif at the exon ends. Only NVS-SM2 shows a clear pattern of GGA motif at the 5’ss, while the other compounds show the inclusion of exons with the canonical nAG motif (Figure 3C, Supplementary Figure 4).

**Figure 4.**
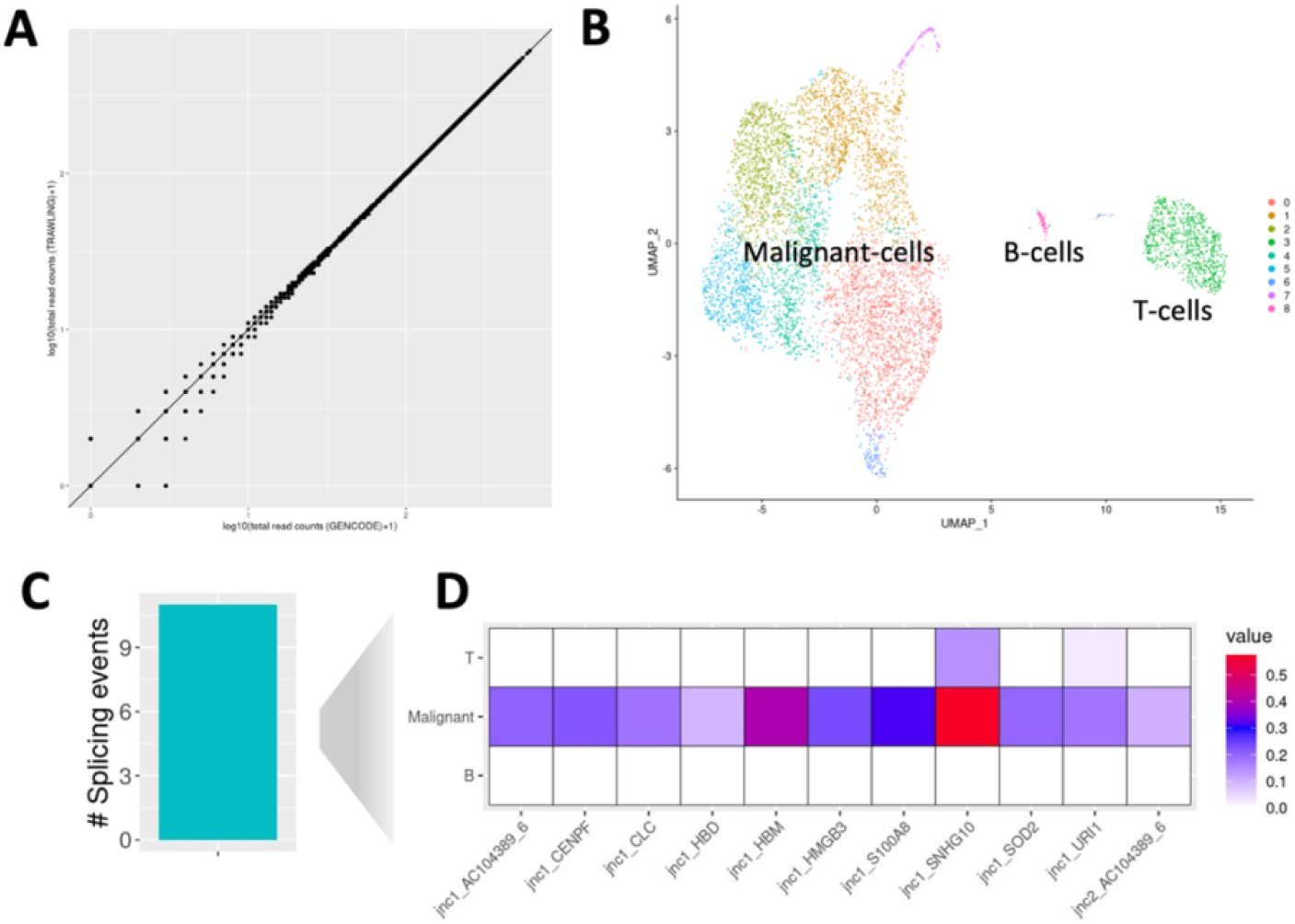
scRNA-seq data analysis. (A) Scatter plot of gene expression levels for 400 randomly selected cells aligned to the last 400 nucleotide of cDNA of each transcript (x-axis) and to TRAWLING (y-axis). (B) UMAP projections of scRNA-seq data. (C) Number of splicing events showing a statistically significant difference between the malignant cells clusters and B/T-cells clusters. (D) Normalized expression value of each differential junction across the clusters.

Importantly, these results also demonstrate TRAWLING’s ability to capture differential splice motif usage from 3’ sequencing alone.

### scRNA-seq data analysis

As in the Drug-Seq dataset, the concordance between the number of reads obtained by aligning to TRAWLING and to the last 400 nucleotides of cDNA in the scRNA-seq dataset was very high (Figure 4A, R^2^ equals to 0.999994).

Then, we conducted a differential splicing analysis between malignant and non-malignant cells (Figure 4B). We identified eleven splicing events showing a statistically significant difference between the Malignant-cells clusters and the B/T-cells clusters (Figure 4C-D). Of those eleven genes, several have previously been shown to play a role in AML, from its initiation to its progression. For instance, the expression of *S100A8* in leukemic cells is a predictor of low survival in AML patients (Nicolas et al., 2011) and its therapeutic targeting has been proposed to improve treatment efficiency in AML. (Mondet et al., 2021). Furthermore Charcot-Leyden crystals (*CLC*) were found in the bone marrow of patients with AML (Khrizman et al., 2010; Radujkovic et al., 2011; Manny et al., 2012). The small nucleolar RNA host gene (*SNHG*) family members have been revealed to be oncogenes in several cancers (Zimta et al., 2020) and *SNHG10* was found to be upregulated in AML patients (Xiao et al., 2021). Also, the authors explored the interaction between *SNHG10* and *miR-621*, suggesting that the former might be targeted by the latter to suppress the proliferation of AML cells. *Hmgb3* RNA has also recently been found to be part of an embryonic stem cell-like transcription signature that was defined in mouse models of mixed-lineage leukemia-mediated leukemic transformation and was also reported to be transiently upregulated during myeloid differentiation (Petit et al., 2010; Somervaille et al., 2009).

### Bulk-RNAseq data analysis

To check whether we were not incorrectly mapping reads when aligning to TRAWLING, the reads were aligned to TRAWLING and to the cDNA of each transcript. As previously shown for the 3’ RNA-seq data analysis, we obtained a high concordance between the two alignment results (Figure 5A, R^2^ equals to 0.999937).

**Figure 5.**
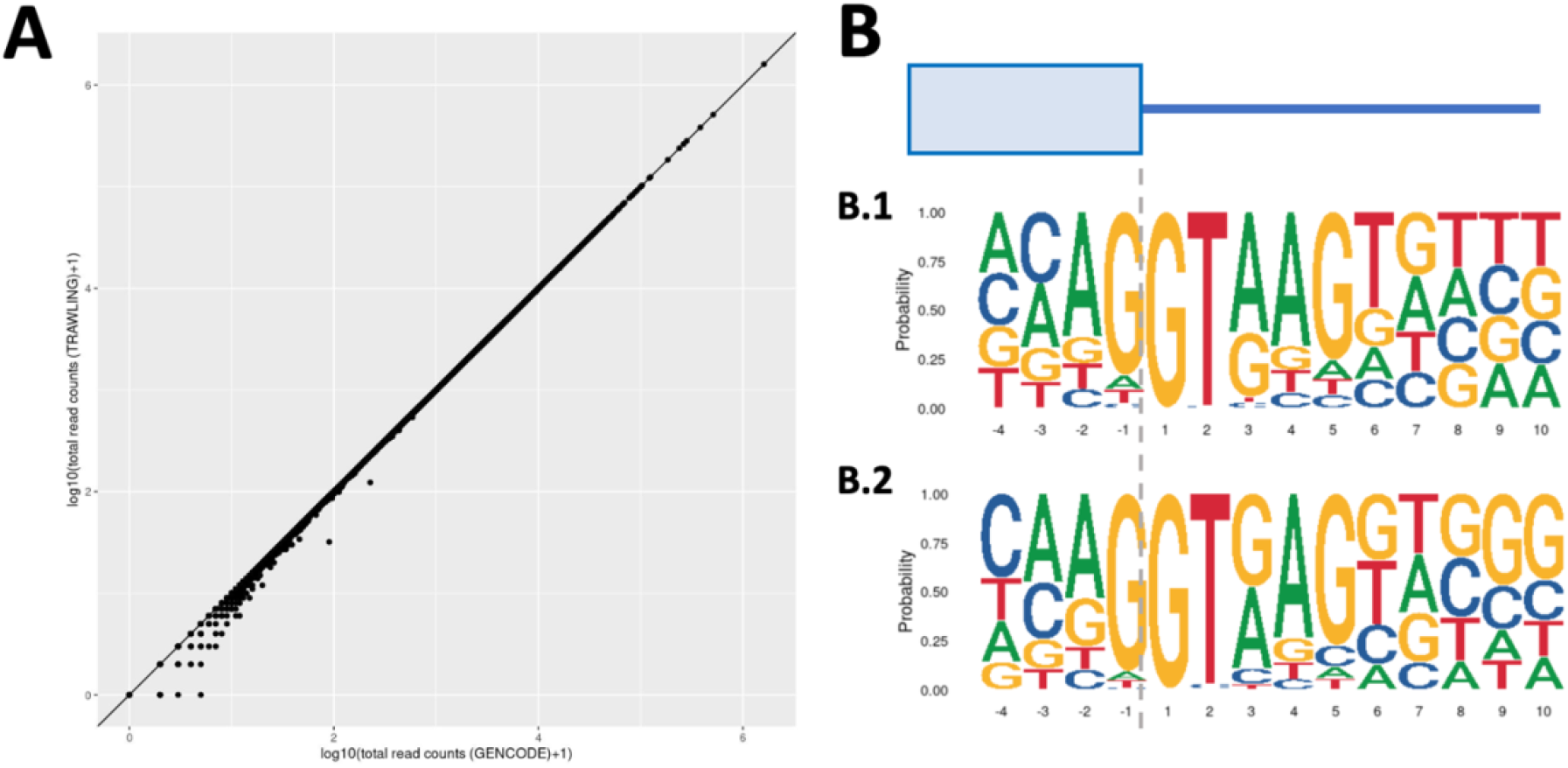
Bulk-RNAseq data analysis. (A) Scatter plot of gene expression levels estimated aligned bulk-RNAseq reads (MUT sample) to the cDNA of each transcript (x-axis) and to TRAWLING (y-axis). (B) Sequence motifs of 5’ splice site of splicing events showing (B.1) a statistically significant difference between WT and the MUT samples (exons skipping events in the latter) and (B.2) a statistically significant difference between MUT and WT samples (exons inclusion events in the former). Among the nucleotide differences between B.1 and B.2 sequence motifs, we can identify the T and G nucleotides preference at the sixth position of the 5’ splice site for B.1 and B.2 respectively, as previously showed (Shuai et al., Nature, 2019)

To evaluate whether the A>C mutation at the third base of U1 snRNA could affects the production of specific aberrant isoforms, we performed a differential splicing analysis using the MUT and the WT samples. We identified several splicing events showing a statistically significant difference between the MUT and the WT. Introns with increased excision were highly enriched in the G at the 6th, 8th, 9th, 10th positions of the 5’ splice site. It suggests that the A>C mutation at the 3 base of U1 could shift the T motif preferably towards a G (Figure 5B). This result is consistent with previous results (Shuai et al., 2019).

## Discussion

In this work, we presented TRAWLING, a transcriptome reference aware of splicing events, that can be built starting just from a gtf annotation file. The performances and the benefits of TRAWLING were evaluated in three different transcriptome datasets: bulk RNA-seq, scRNA-seq and DRUG-seq.

Compared to aligning transcriptome data to just the cDNA of each transcript, TRAWLING enabled the detection of splicing events in a fast, simple and accurate way. In addition, TRAWLING did not misalign or lose reads, which demonstrates that it can be used by default without loss of generality for gene level quantification. Upon further development, transcript level quantification would also be possible.

In addition, TRAWLING can aggregate read counts based on the donor and acceptor splice motifs and the advantages of this feature have been shown in the context of DRUG-seq and bulk RNA-seq data analysis. In the scRNA-seq data analysis, TRAWLING enabled the identification of different splicing events relevant to AML genes (e.g. *S100A8, CLC*).

Together, these 3 dataset analyses demonstrate that TRAWLING is widely applicable to transcriptomic data of different types and enables flexible downstream analysis.

Interestingly, the approach developed to build TRAWLING could easily be applied to extend the reference transcriptome to non-canonical splicing events (Supplementary Figure 5-6). This is an important aspect since the available transcript reference annotations are meant to capture the canonical isoforms rather than those symptomatic of very rare or diseased states. Non-canonical splicing events can potentially lead to disease-causing aberrant transcripts but can also offer new therapeutic strategies (Sibley et al. 2016; Wang et al. 2021). It is estimated that DNA variants in non-canonical splice motif are underrepresented in existing public databases, despite conveying a similar risk of disease as other DNA variants (Lord et al., Genome research 2019). Non-canonical splicing events could be collected from publicly available resources such as Snaptron (Wilks et al., Bioinformatics, 2018), a database containing exon-exon junctions from thousands of RNA-Seq samples (Supplementary Figure 7). They could also be collected from published studies, like in Wang et al. 2021, in which the authors detected and annotated a specific type of non-canonical splicing event, called exitron (cryptic introns with both splice sites inside an annotated exon), across 33 cancer tissues.

In summary, we believe that TRAWLING is suitable for various splicing studies and will provide new insights, such as the discovery of unannotated drug mechanism of actions and new potential targets. We believe future expansions of TRAWLING will further increase its impact on transcriptomic data analysis.

