## Supplementary Material for "TRAWLING: a Transcriptome Reference Aware of spLIciNG events"

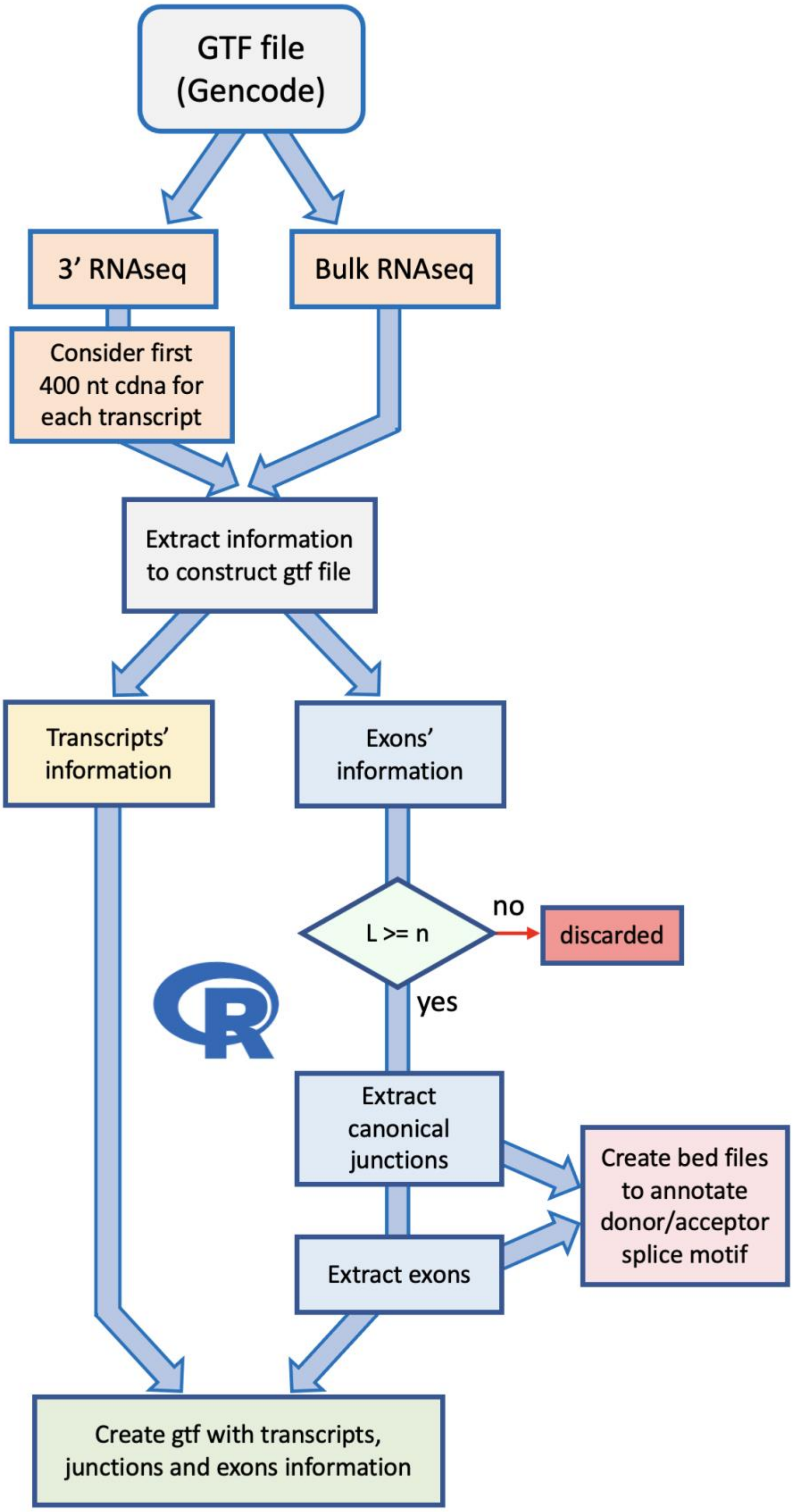


**Figure 1 Create GTF and bed files for TRAWLING.** Starting from the gtf annotation file, the exons, junctions, and cDNA information are extracted and annotated in a new gtf file. The donor and acceptor splice motifs are annotated for every exon and junction.

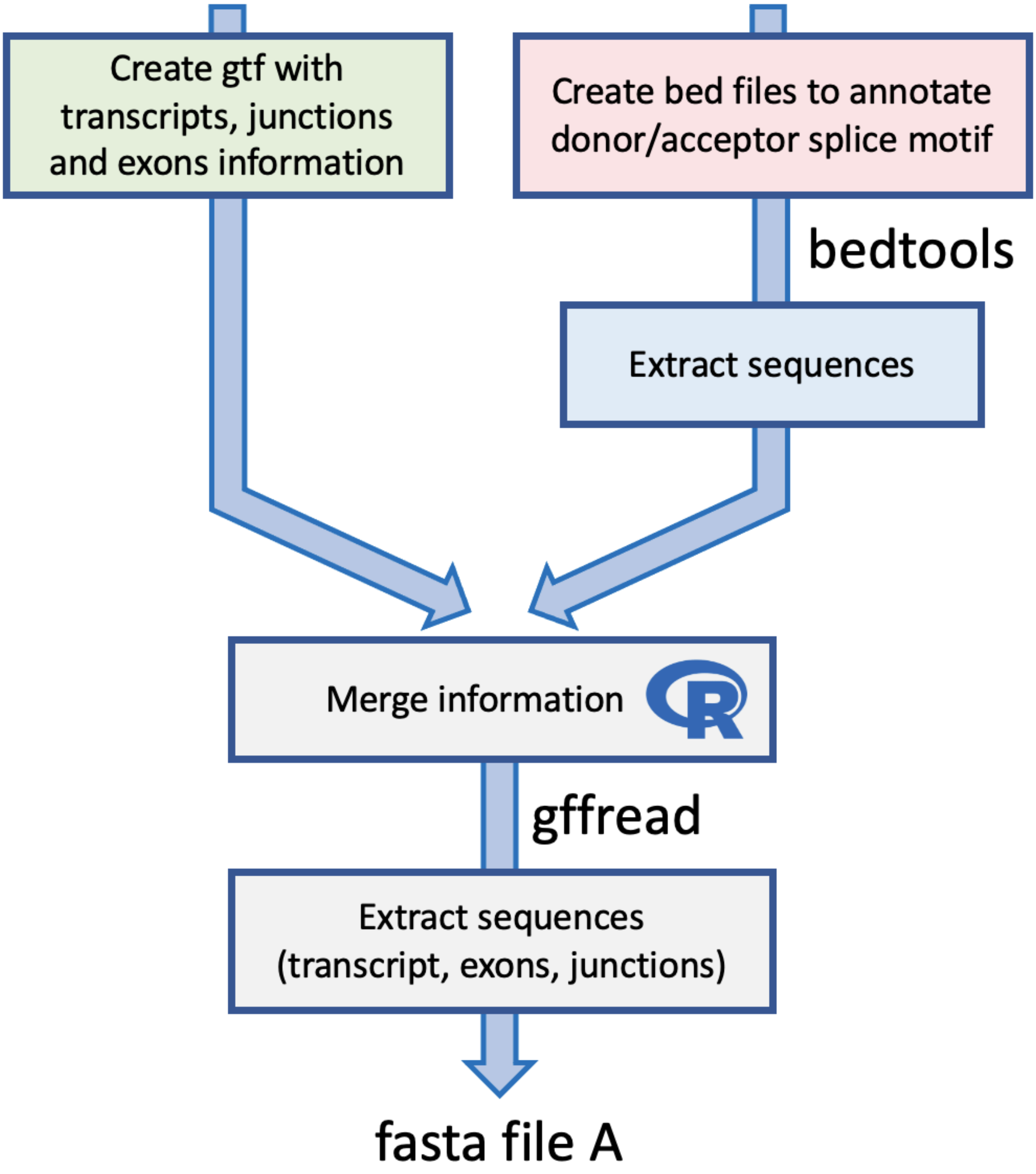


**Figure 2 Build TRAWLING from the GTF file.** The information included in the gtf file format and the donor/acceptor splice sequences are merged into a new gtf file. The transcript reference fasta is obtained using TRAWLING's gtf file and gffread tool (Pertea et al., 2020).

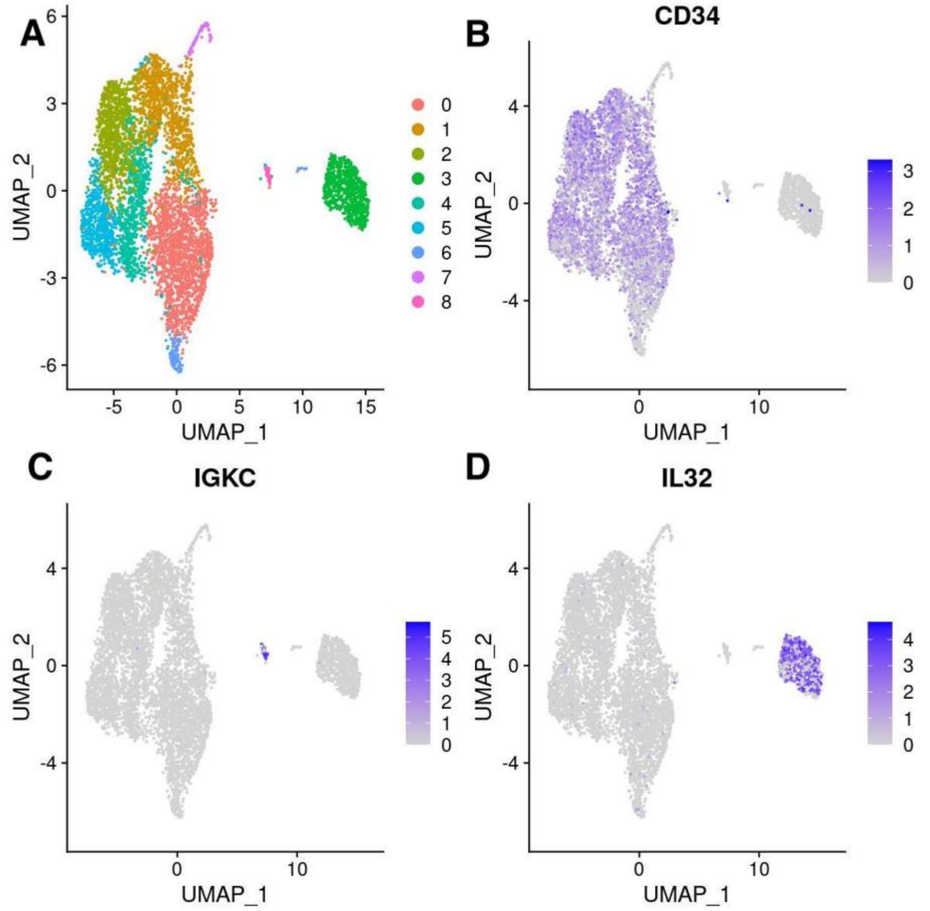


**Figure 3 Analysis of AML scRNAseq dataset** (A) UMAP representation (B)-(D) Visualization of markers' expression across clusters. (B) CD34 is a leukemic stem cell marker in AML. (C) IGKC is a marker gene for B-cells (Schmidt et al., 2012). (D) IL32 is a marker gene for T-cells (Dahl et al., 1992). The plot is generated using a publicly available scRNAseq dataset (GEO accession GSM4317810) presented in Johnston et al. (2020).

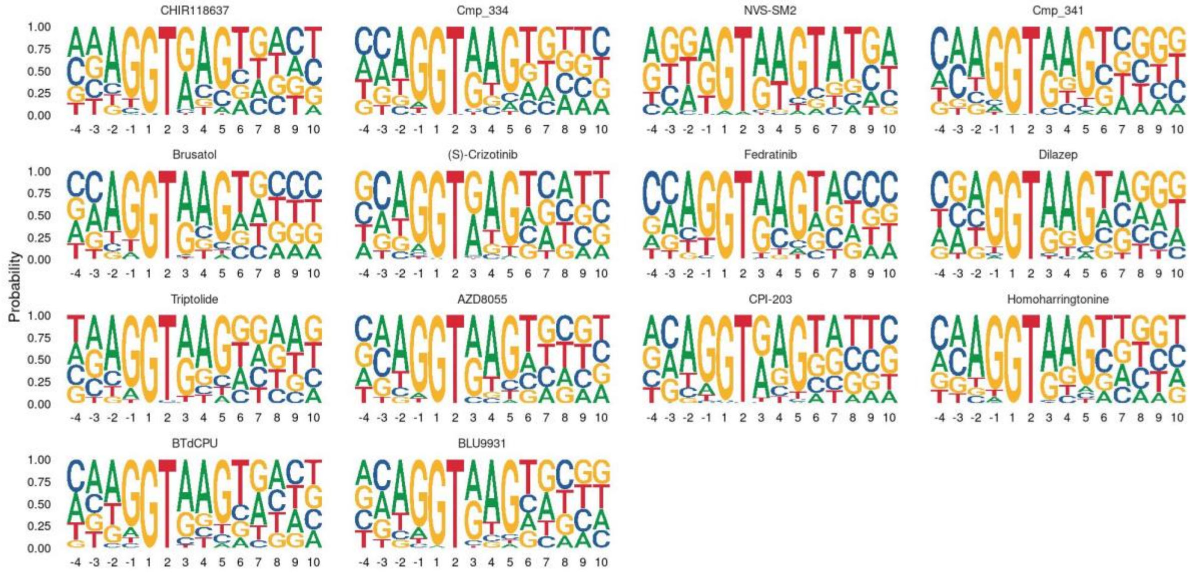


**Figure 4 Donor splice motifs enrichment of each compound included in the DRUG-seq dataset**. For each compound,the splicing signal motifs at 5'ss is shown. The plot is generated using a publicly available DRUG-seq dataset (GEO accession GSM5357052) and includes only splicing events that showing a statistically significant difference between DMSO and the considered compounds (title name of each motif plot), when used at a dosage of 10uM.

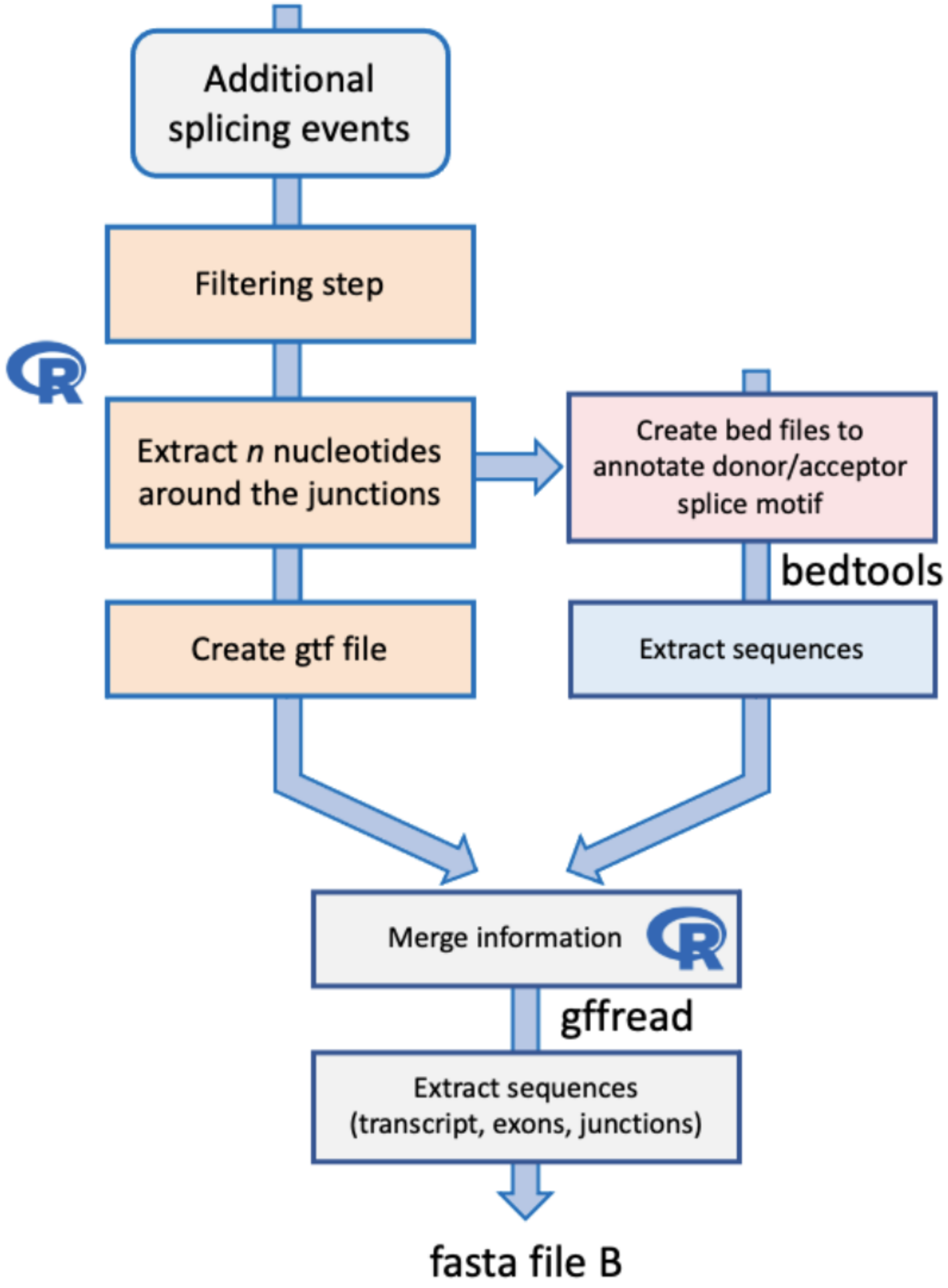


**Figure 5 Including additional splicing events in TRAWLING (**e.g. EXITRON events, non-canonical splicing events collected from SNAPTRON (Wilks et al., Bionformatics, 2018)).

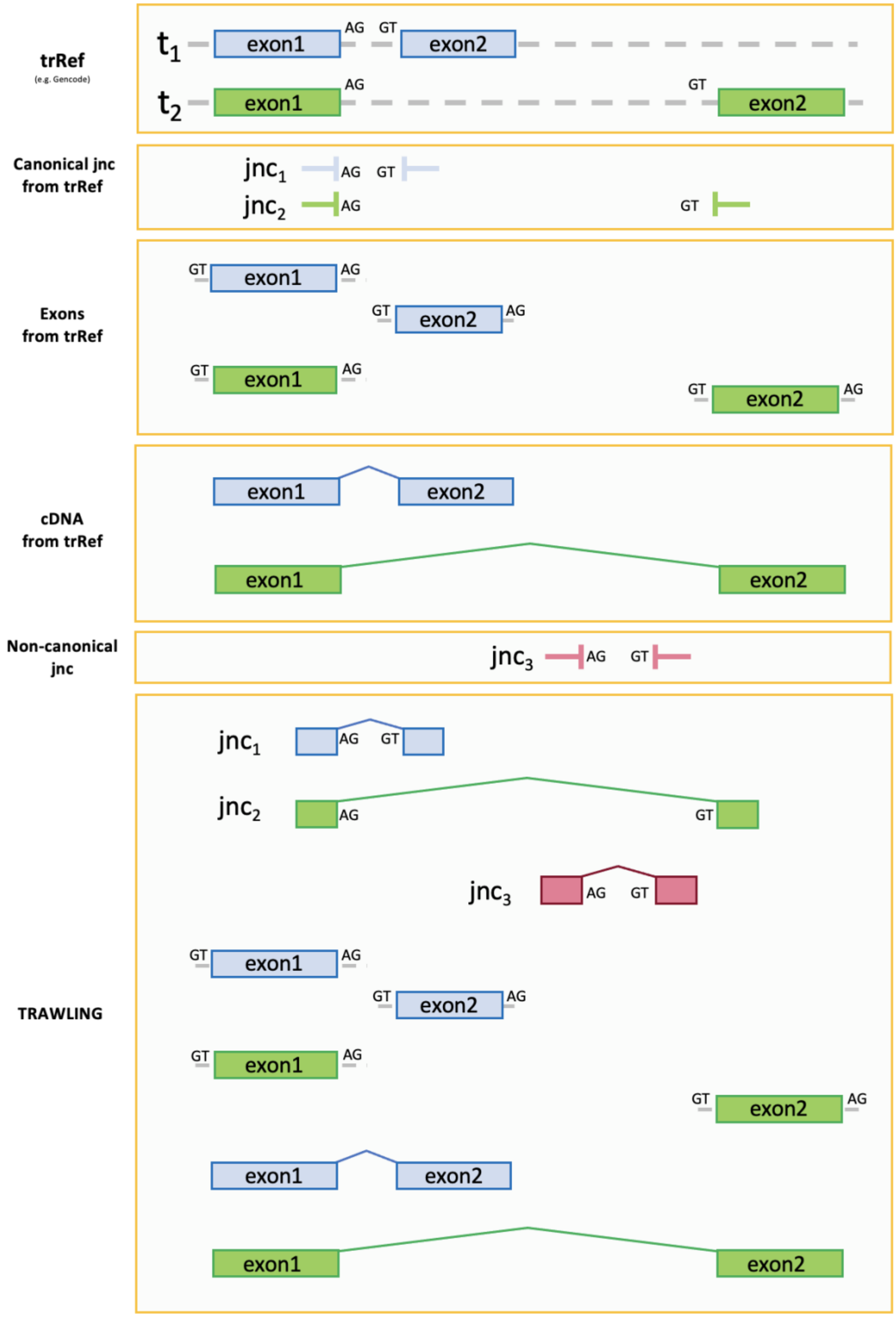


**Figure 6 Extend TRAWLING by including non-canonical splicing events.** In this conceptual example, the information of unannotated splicing events is collected and included in TRAWLING. The non-canonical splicing events can be retrieved from source such as Snaptron (Wilk et al., 2018) or publicly available studies (e.g. Want et al, 2021)).

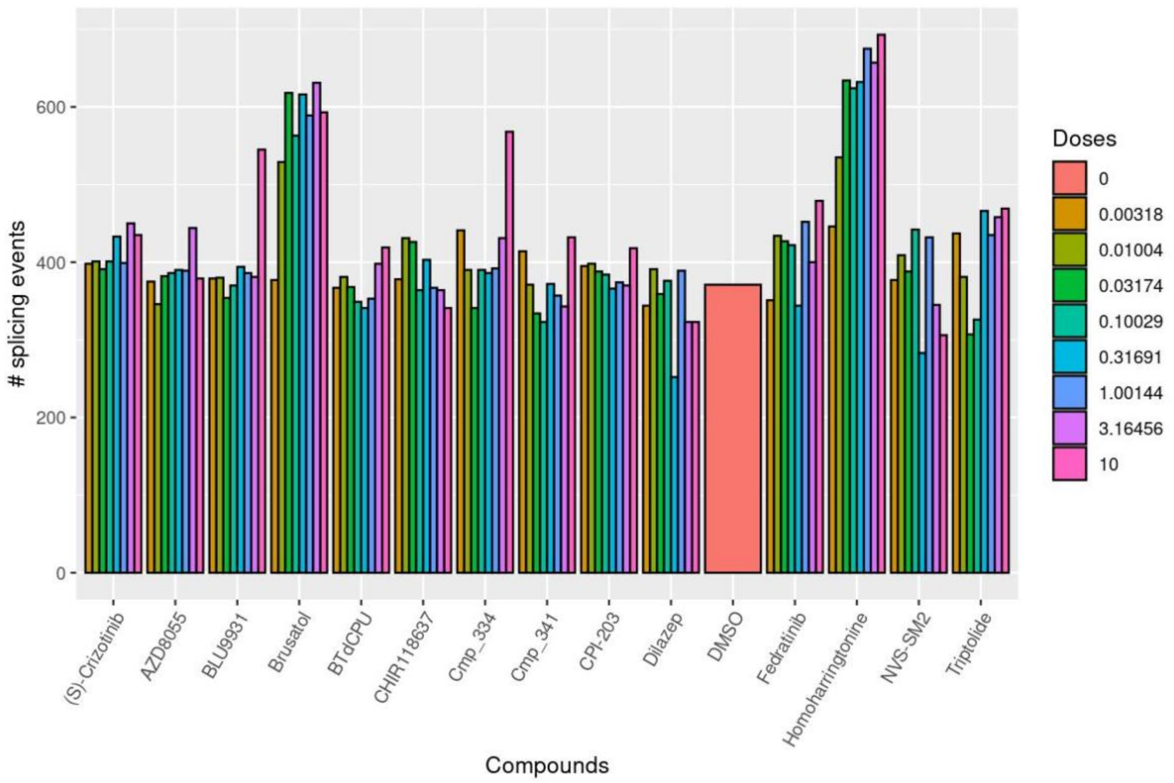


**Figure 7 Non-canonical splicing events detect in DRUG-seq data.** The non-canonical splicing events were collected from Snaptron (Wilk et al., 2018) and included in TRAWLING. The plot shows the number of non-canonical splicing events detected in the DRUG-seq dataset. This result is generated using a publicly available DRUG-seq dataset (GEO accession GSM5357052) presented in Li J et al., (2021).
